# Degradation of lactoferrin caused by droplet atomization process via two-fluid nozzle: The detrimental effect of air–water interfaces

**DOI:** 10.1101/2021.12.06.471411

**Authors:** Huy M. Dao, Sawittree Sahakijpijarn, Robert R. Chrostowski, Chaeho Moon, Filippo Mangolini, Zhengrong Cui, Robert O. Williams

## Abstract

Biological macromolecules, especially therapeutic proteins, are delicate and highly sensitive to degradation from stresses encountered during the manufacture of dosage forms. Thin-film freeze-drying (TFFD) and spray freeze-drying (SFD) are two processes used to convert liquid forms of protein into dry powders. In the production of inhalable dry powders that contain proteins, these potential stressors fall into three categories based on their occurrence during the primary steps of the process: (1) droplet formation (e.g., the mechanism of droplet formation, including spray atomization), (2) freezing, and (3) frozen water removal (e.g., sublimation). This study compares the droplet formation mechanism used in TFFD and SFD by investigating the effects of spraying on the stability of proteins, using lactoferrin as a model. This study considers various perspectives on the degradation (e.g., conformation) of lactoferrin after subjecting the protein solution to the atomization process using a pneumatic two-fluid nozzle (employed in SFD) or a low-shear drop application through the nozzle. The surface activity of lactoferrin was examined to explore the interfacial adsorption tendency, diffusion, and denaturation process. Subsequently, this study also investigates the secondary and tertiary structure of lactoferrin, the quantification of monomers, oligomers, and ultimately, aggregates. The spraying process affected the tertiary structure more negatively than the tightly woven secondary structure, resulting in a 1.5 nm red shift in peak position corresponding to the Tryptophan (Trp) residues. This conformational change can either (a) be reversed at low concentrations via relaxation or (b) proceed to form irreversible aggregates at higher concentrations. Interestingly, when the sample was allowed to progress into micron-sized aggregates, such a dramatic change was not detected using methods such as size-exclusion chromatography, polyacrylamide gel electrophoresis, and dynamic light scattering at 173°. A more complete understanding of the heterogeneous protein sample was achieved only through a combination of 173° and 13° backward and forward scattering, a combination of derived count rate measurements, and micro-flow imaging (MFI). Finally, compared to the low-shear dripping used in the TFFD process, lactoferrin underwent a relatively fast conformational change upon exposure to the high air-water interface of the two-fluid atomization nozzle used in the SFD process as compared to the low shear dripping used in the TFFD process. The interfacial induced denaturation that occurred during spraying was governed primarily by the size of the atomized droplets, regardless of the duration of exposure to air.

## INTRODUCTION

Medicines based on biological macromolecules (e.g., peptides, proteins) are quickly becoming important assets in our fight against disease, alongside conventional synthetically derived small molecules. Macromolecules offer several advantages over small-molecule therapies because they (a) allow for more specificity, which improves their safety profile, and (b) bioengineering processes can tailor macromolecules to keep pace with the evolution of diseases.^1^ However, the extremely complex nature of these macromolecules makes it difficult to understand their structure and their sensitivity to environmental factors, which increases their tendency for denaturation.

With protein-based therapies (e.g., enzymes, therapeutic proteins, polypeptides, antibodies), their native or folded structures and the delicate spatial arrangement of their polypeptide chains affects their specific therapeutic efficacy. The native form, in terms of the energy landscape, is usually the most stable form, possessing lower Gibbs free energy than other forms.^3^ However, the difference in free energy between the native, partially unfolded, and completely unfolded forms can be relatively small in many cases, ranging from 5–20 kcal/mol.^3^ As a result, minor changes in environmental factors (e.g., pH, ionic strength, mechanical energy input), or even exposure to the air–water interface, can result in the unfolding of the protein.^4–8^ Due to their exposed hydrophobic pockets, denatured proteins can self-assemble, resulting in protein aggregates that can either decrease their therapeutic efficacy or produce unwanted immunogenicity.^1^

With the rapid spread of the SARS-CoV-2 virus, the need for the mass vaccination of entire populations around the world has created an urgent public health need for greatly improved cold-chain storage.^9^ Ideally, cold-chain storage must be capable of maintaining the biological activity of a protein-based vaccine or therapeutic over an extended period of time at storage temperatures that can be achieved in all countries. This is critical for countries that lack the infrastructure needed to support the storage and distribution of protein-based vaccines or therapeutics without compromising their stability. Moreover, the effective delivery of biological macromolecules to the lungs is a challenging but important task because it increases efficacy and reduces side effects in the treatment of pulmonary diseases, including viral infections like COVID-19.^10^

In light of this, dry powder inhalation is an excellent choice because it addresses two challenges at once: (a) delivering the macromolecular drugs directly to the site of therapeutic action and (b) increasing the storage stability of the macromolecular drug (e.g., protein-based therapeutic).^11,12^ The conversion of the liquid form of the protein (which readily causes various types of denaturation, ranging from chemical to physical) into its solid form significantly extends the shelf-life stability of the formulation without the need for storage at extremely low storage temperatures. For example, the mRNA–lipid nanoparticle vaccines for COVID-19 currently approved in the United States require temperatures as low as −20 °C or even −80 °C.^9^ Conventional lyophilization, which employs slower freezing on a shelf as well as sublimation, has been used extensively with proteins for this purpose. However, its applicability to more advanced macromolecule formulations (e.g., formulations that contain lipid nanoparticles to facilitate the delivery of the mRNA) has not been sufficiently demonstrated.^13^ Additionally, conventional lyophilization alone does not create an inhalable dry powder form due to the resulting lyophilized cake’s high rigidity.^14^ Hence, the preparation of dry powders suitable for inhalation requires either an upstream technology or a downstream processing technique to convert the rigid, bulky lyophilized cake into an inhalable powder.

In this regard, tTwo engineering techniques for preparing aerosolizable dry powder formulations suitable for pulmonary delivery have been reported: *spray freeze-drying* (SFD)^5,15,16^ and *thin-film freeze-drying* (TFFD)^17^ (Fig. 1). While SFD terminology has been used to describe a heterogeneous set of techniques, it typically involves the formation of small, micron-sized droplets produced by spray-atomizing a liquid through a two-fluid, high-pressure nozzle directly into a cryogen or cryogenically cooled environment. This frozen slurry of particles is collected, and ice is removed via sublimation through lyophilization. The TFFD process involves the formation of large millimeter-sized drops produced using a low-shear nozzle. These drops are applied to a cryogenically cooled surface (e.g., the stainless steel surface of a rotating drum apparatus).^18^ It is evident that protein stability is affected by differences in the mechanism of droplet formation and subsequent freezing.

**Figure 1.**
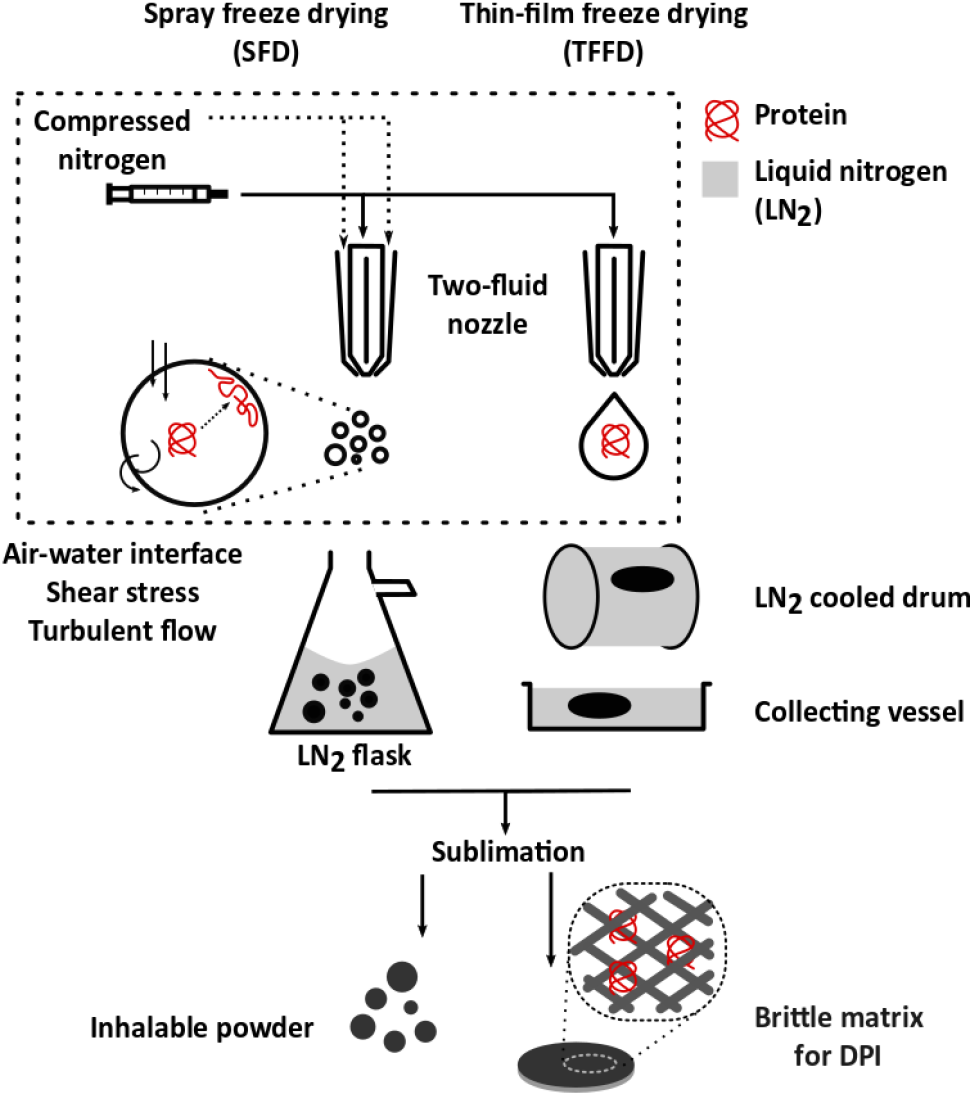
Schematic diagram of two powder engineering techniques suitable for pulmonary delivery: spray freeze-drying (SFD) and thin-film freeze-drying (TFFD). The rectangular selection denotes the focus of this study.

The resultant powders formed using SFD and TFFD are highly porous with a low mass median aerodynamic diameter (MMAD) capable of depositing in the lower region of the lung.^14,19–51^ Although it is capable of producing inhalable powders, SFD has several inherent drawbacks. These drawbacks include the formation of a very large air–water interface (or surface-to-volume ratio) and the use of high-shear stress on the liquid as it is forced through the two-fluid nozzle accompanied by turbulent flow.^15^

SFD can be acceptable for preparing dosage forms that contain small molecules, but in order to avoid protein denaturation at the air–water interfaces, mitigation approaches must be considered when translating the SFD process into the formulation of macromolecules.^22^ In these aspects, TFFD mitigates these drawbacks by requiring a lower-stress procedure that includes applying comparatively larger protein liquid drops (~4–7 mm) onto a cryogenically cooled stainless steel rotating drum. After this, the formed frozen thin films are collected and subsequently sublimated to remove the frozen solvent (e.g., water). TFFD produces dry powders for inhalation suitable for many small molecules, including remdesivir,^23^ tacrolimus,^18^ niclosamide,^24^ voriconazole,^20,25^ and a few macromolecules such as lysozyme,^17^ lactate dehydrogenase,^16^ ovalbumin-adsorbed vaccine adjuvant,^11^ and the norovirus vaccine.^26^ The TFFD-processed powders have excellent fine-particle fractions (FPF ~70–90%) and median mass aerodynamic diameters (MMAD < 2–5 μm) for dry powder inhalation.^11,25,27^

A more in-depth understanding of the effects of stress applied to the macromolecule during droplet formation is needed to expand the application of SFD and TFFD to the manufacture of inhalable dry powders that contain stable proteins.^28,29^ It was initially assumed that proteins experienced denaturation due to applied shear stress and exposure to the air–water interface. However, some studies point out that proteins can still maintain their tertiary structure even under extreme shear stress (up to 10^7^ s^−1^ without the presence of an interfacial area).^30,31^ As a result, the primary detrimental factors encountered with spray-atomizing techniques are believed to be exposure to a high surface area-to-volume ratio and turbulent flow.

Lactoferrin was chosen as the model protein, with the long-term vision of preparing inhalable dry powder formulations suitable for lung delivery. In terms of therapeutic justification, lactoferrin is an 80 kDa protein possessing activities against airborne viral pathogen–related diseases (e.g., COVID19) via three main mechanisms: (a) blocking virus attachment to epithelial cells, (b) enhancing immune responses, and (c) regulating the cytokine storm.^32–39^ Lactoferrin surface unfolding has been previously studied in the context of the tear film, and deep molecular conformation has been studied via small-angle neutron scattering.^40^ This study examined the initial steps of the SFD and TFFD techniques related to the formation of smaller droplets and larger drops to be frozen, and it compared their respective mechanisms of droplet/drop formation (Fig. 1). The liquid protein formulation was not frozen or sublimated, so the stress factors relevant to the air–water-induced denaturation of the protein could be isolated.

## EXPERIMENTAL SECTION

### Materials

Lactoferrin was purchased from Lactoferrin Co. (Melbourne, Australia). Polysorbate 20, sodium lauryl sulfate, acetonitrile, methanol (HPLC grade), trifluoroacetic acid, phosphate buffer saline, and BCA protein assay were purchased from Fisher Scientific (Pittsburgh, PA, USA).

### Spraying process

Lactoferrin was dissolved in water or in a 0.01 M PBS pH 7.4 solution, then centrifuged at 4,000 rpm at 4 °C for 30 min. The supernatant was filtered through a 0.45 μm PES membrane to eliminate insoluble inorganic particles. The lactoferrin solution was sprayed using a two-fluid pneumatic nozzle with a 0.7 mm nozzle diameter and a 1.5 mm nozzle cap. Filtered dried nitrogen gas was used as the atomizing gas at a flow rate of 0–40 L/min, controlled via a critical flow controller (TPK 2000, Copley Scientific, Nottingham, UK) positioned at the nozzle input. The feed rate was controlled using a peristatic pump. The solution was sprayed into 50 mL glass vials and stored at 4–8 °C until they were analyzed.

### Analysis of lactoferrin structure

Lactoferrin was examined for changes in its secondary and tertiary structures using circular dichroism (CD) spectroscopy. CD spectra were collected using a JASCO CD180 spectrophotometer from 190–260 nm for a near-UV study of the secondary structure and from 250–350 nm for a far-UV study of the tertiary structure. Lactoferrin solutions were prepared at concentrations of 300 μg/mL and 1,000 μg/mL for the analysis of the secondary and tertiary structures, respectively. All spectra were acquired at 20 °C in a 5 mM phosphate buffer at pH 7.4 using quartz cuvettes with a 1 mm or 10 mm light path. All recorded spectra were the result of five scans conducted at a scanning speed of 50 nm/min. A blank spectrum was acquired from the corresponding media and subtracted accordingly. All spectra were normalized for their lactoferrin concentration, which was obtained using the BCA protein assay or their absorption at 280 nm from the UV spectra.

### Analysis of aggregation

A Zetasizer ZS (Malvern Analytical, UK) was used for the dynamic light scattering (DLS) measurements to monitor the aggregation tendency of the proteins. A lactoferrin solution was loaded into a semi-micro disposable plastic cuvette and sealed with parafilm. The air space of the cuvette containing the sample was flushed with filtered nitrogen gas for 1 min before sealing to minimize chemical oxidation reactions.

The DLS measurements were carried out at 173° and 13° scattering angles to monitor the smaller and larger protein particle populations, respectively. The derived count rate, which is measured in *kilo-count-per-second* (kcps), is a normalized parameter obtained by the normalization of the count rate using the Zetasizer attenuator setting. The derived count rate is a nonlinear function of both particle size and the number of particles presented in the system. It is collected and reported as a parameter to track protein aggregation. All measurements were performed in triplicate.

### Size exclusion chromatography (SEC)

SEC was utilized to analyze the monomer and oligomer content of the sprayed lactoferrin sample. SEC was performed using an Agilent AdvanceBio SEC 300 A 2.7 μm column, with a mobile phase consisting of 0.2 M phosphate buffer at pH 6.8 and 0.3 M sodium chloride at 0.35 mL/min. The detector was set at 220 nm and 280 nm with a 5 μL injection volume. The primary monomer peak of lactoferrin was eluted at 4.1 min.

### Protein quantification

The lactoferrin content was determined using a BCA pierce assay (Thermo Scientific). This process is based on the manufacturer’s protocol, in which (1) 100 μL of lactoferrin-containing samples were incubated with 2.0 mL of working reagent at 37 °C for 30 min and (2) the absorbance at 562 nm was measured. Results from the BCA assay were used to normalize the protein concentration when necessary.

### Surface tension measurement

The interfacial tension, or surface pressure, was measured using a Theta Flex Plus tensiometer, which is a drop shape analysis system (Biolin Scientific). The lateral pixel recorded by the instrument was calibrated using a precise 4 mm diameter steel ball, and the interfacial tension of the pure water solution was measured at approximately 72 mN/m to confirm the calibration of the camera. The droplet shape of the lactoferrin solution was monitored automatically to track changes in the interfacial tension over a period of 200 s. A droplet with a volume of ~10 μL was dispensed at 10 μL/s via automated pipet. Evaporation compensation was turned off to avoid the addition of newly native proteins to the system. In this case, the evaporation of the water is negligible (< 0.2 μL) owing to the short duration of the experiments. The dynamic surface tension data were fitted to the Hua–Rosen model using IgorPro (v.9, WaveMetrics Inc., USA).

### Micro-Flow Imaging

The lactoferrin samples were characterized for subvisible aggregates using micro-flow imaging (MFI) MFI5100 (ProteinSimple, San Jose, CA) equipped with a Bot1 autosampler. Specifically, 0.9 mL of lactoferrin protein solution was analyzed at a flow rate of 0.17 mL/min with a 300 μm flow cell. Before each sample is analyzed, the flow cell was flushed multiple times with particle-free water, which contained < 500 particles/mL with an equivalent circular diameter (ECD) ranging from 1 μm to 400 μm. The lactoferrin sample was stirred three times with Bot1 pipette at the lowest possible stirring rate to avoid the formation of stable foam. An MFI view system suit (MVSS) and its built-in image analysis software (ProteinSimple) were used to process the data for the lactoferrin samples that were spray-atomized at various airflow rates. Extrinsic particles (e.g., rubber shards, foreign dust particles) were removed from the data using the “find like particles” feature. Intrinsic particles (e.g., air microbubbles, silicon oil droplets) were removed from the data by applying a filter that counts only particles with an aspect ratio less than 0.9 and a circularity less than 0.9. The data were represented using heat maps that include the aspect ratio, circularity, and ECD of protein particles.

## RESULTS

This study investigated the effect of air–water interface exposure on the structural integrity of the protein lactoferrin. First, the surface-active properties of the protein were studied using a drop-shape analysis tensiometer. Next, the secondary structure and tertiary structure were studied using far-UV and near-UV CD spectroscopy, respectively. Dimer and oligomer content was characterized using SEC. Subsequently, the macroscopic aggregation properties of the protein were investigated using a modified approach with DLS.

First, lactoferrin was examined for its interfacial activity, and it was found to be strongly surface active. As shown in Fig. 2A–B, lactoferrin preferentially accumulates at the air–water interface, where it significantly reduces the surface tension of the air–water interface (from 72 mN/m to ~ 60 mN/m in water). This effect is even more pronounced in a 0.01 M PBS buffer at pH 7.4, where phosphate ions and sodium chloride are presented, and the pH is higher (Fig. 2B). Lactoferrin is a positively charged protein with an abundance of arginine and lysine amino acid residues in its structure. The higher pH and ionic strength likely screened the electrostatic repulsion between adjacent molecules, enabling the protein to pack more tightly at the interface, resulting in lower interfacial tension.^41^ The phosphate ions can also act as a physical crosslinking agent to form a more rigid protein film at the interface, thus resisting ambient vibration.^42^ As a result, the relative amount of noise or fluctuation in the signal was much lower in the PBS samples than in water samples with the same protein concentration (Fig.2B).^41^

**Figure 2.**
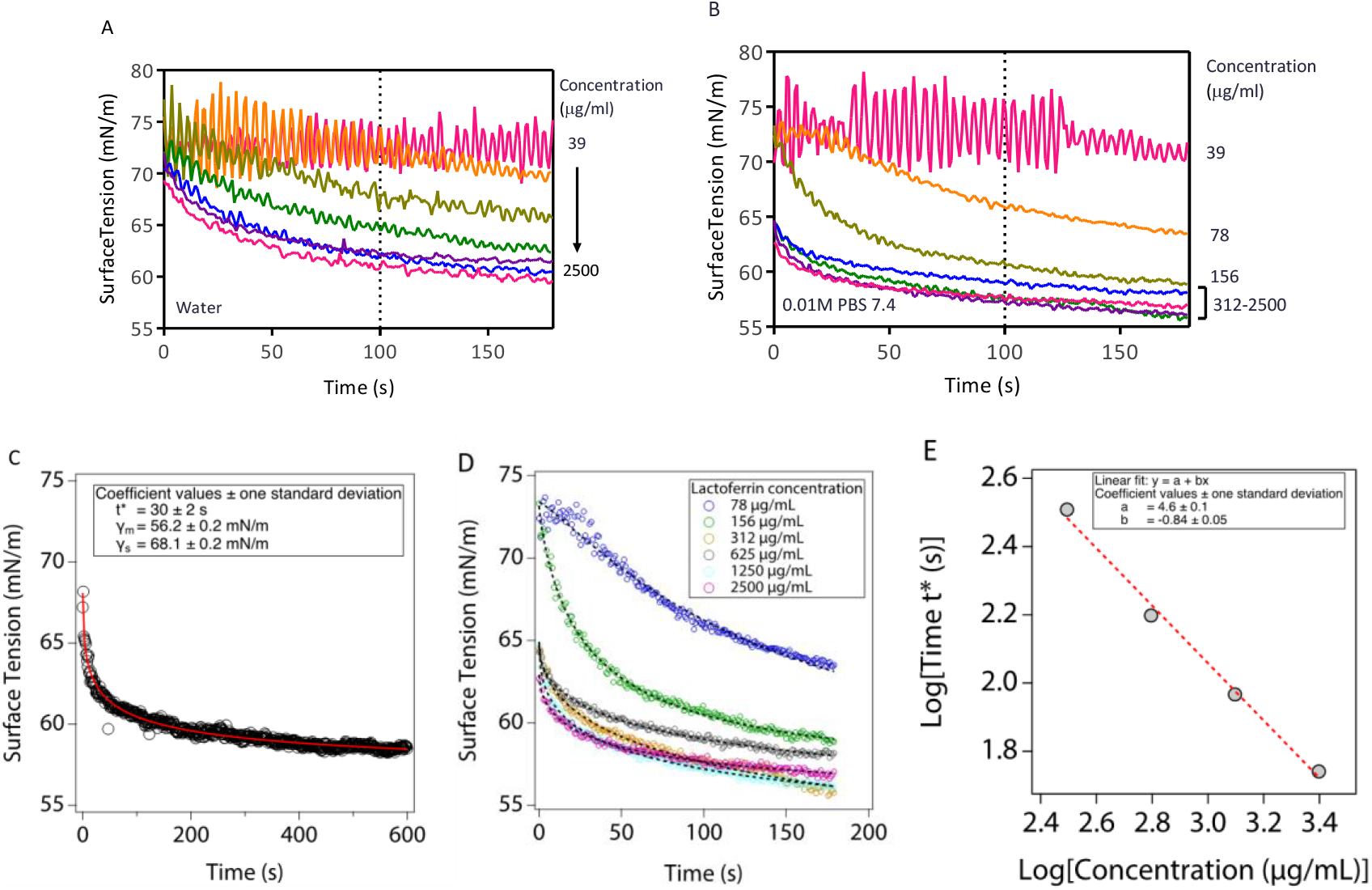
Surface tension of the air–water interface of lactoferrin solutions in water (A) and in 0.01 M PBS at pH 7.4 (B). The representative fitting coefficient and fitting curve to the Hua–Rosen model to obtain the *t\** value of the lactoferrin solution at 1,000 μg/mL (C) and from 78–2,500 μg/mL(D). The concentration dependent on log *t*\*(E).

To investigate the interfacial activity of lactoferrin, the protein solution was dispensed at 10 μL/s into ~10 μL droplets, resulting in a < 1 s preparation time before recording the measurements. Notably, the surface tension of the PBS samples with protein concentrations > 156 μg/mL began at a value different from 72 mN/m, which is the surface tension of pure water (Fig. 2B). The deviation from this surface tension value suggests that the air–water interface of those samples had already been covered by at least 50% of the close-packed monolayer of lactoferrin from the time data collection began.^43^ On the other hand, the surface tension value for all lactoferrin–water solutions converged at about 72 mN/m at the early stage of the measurements. This suggests that either the 50% coverage had not yet been attained or the monolayer was loosely packed.^43^ Nonetheless, these results demonstrate that lactoferrin is a surface-active protein with an equilibrium surface tension on par with several monoclonal antibodies, hence it is susceptible to air–water interfacial denaturation.^22,43^

The dynamic surface tension characteristics of lactoferrin were further investigated via fitting the data with the Hua–Rosen model^44^ to understand its adsorption kinetics, using the equation

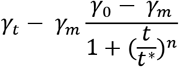

Specifically, γ_0_ represents the surface tension of the pure solvent, γ_t_ is the surface tension of the solution at time *t*, and γ_m_ is the surface tension of the solution at the meso-equilibrium region. The variable *t^*^* represents the time required for the surface tension to reach the midpoint between γ_0_ and γ_m_, which is an indicator of the rate of adsorption kinetics. Fig. S1 shows a typical curve of the change in dynamic surface tension with time. The Hua–Rosen model proposes to divide the adsorption curve into four regions: *induction, rapid fall, meso-equilibrium*, and *equilibrium*. Fig. 1C and Fig. S2 show the representative fitting coefficient and the fitting curve of the dynamic adsorption of the lactoferrin solution. The calculated characteristic time *t*\* (the time at which the surface tension is halfway between the value of the pure solvent and the value at meso-equilibrium) is 30 ± 2 s. These results indicate that the adsorption process of lactoferrin at the air–water interface is much slower than the kinetics of denaturation, or unfolding, of the protein, which is expected to occur within a nanosecond time range.^4^ The protein diffusion process is also much slower than the droplets’ travel time from the nozzle tip to the container wall, which happens within a few milliseconds. Thus, it is assumed that lactoferrin diffusion is responsible for only a negligible portion of the overall protein denaturation that occurs during spray atomization.

The Ward and Todai model^45^ assumes the adsorption process is controlled only by the bulk diffusion of surface-active molecules. It predicts that the time required to attain a given surface coverage is proportional to (1/C_b_)^2^, where C_b_ is the bulk concentration of the adsorbate. After fitting the experimental log (*t*\*) versus log (C_b_) with a linear function, the computed slope is −0.84 ± 0.05 (Fig. 2D–E). This value differs from the hypothetical value of −2.00, where the process is purely diffusion controlled.^45^ The deviation of the slope from the value predicted by the Ward and Todai model can be explained by the fact that diffusion is not the only mechanism that controls the adsorption kinetics of surface-active molecules of a particular size and structure. The exchange kinetics of surface-active molecules (e.g., lactoferrin) between the surface and the bulk liquid adjacent to the surface must also be taken into consideration. The process of matter exchange was reported to require protein molecules to overcome an adsorption energy barrier. The whole process of lactoferrin adsorption follows the so-called “mixed adsorption kinetics.”^46^

After the surface activity of lactoferrin was confirmed, the potential denaturation by adsorption at the air–water interface was studied by exposing the protein to a large interfacial surface area created via spray-atomizing techniques. Specifically, the lactoferrin solution was sprayed into low-micron-sized droplets with diameters in the range of 20–200 μm using a two-fluid nozzle and compressed nitrogen gas. This droplet size range is reported for the SFD process using a two-fluid nozzle.^15,36^ The atomizing airflow rates in this study range from 0 L/ min, which represents the drop-forming conditions used in TFFD; and 10–40 L/min, which represents the conditions used in atomized spray-related techniques in particle formation technologies such as SFD, spray-drying, and spray freezing into liquid (SFL) (Fig. 1).

Fig. 3A–B shows the far-UV CD spectra of the freshly prepared lactoferrin reference samples. The reference spectra were composed of a broad negative band center at 208–210 nm. The shoulder at 215–217 nm suggests a mixture of α-helix and β-sheet conformation (21% α-helix, 31% β-sheet, 12% β-turn, and 36% random coil).^47^ The CD spectra in Fig. 2 conform to the standard spectra of native bovine lactoferrin with only minor variations in peak positions and ellipticities. After heating at 60 °C for 10 min, it was evident that the secondary structure of lactoferrin was significantly disrupted, and its fraction of random coil structure increased. This was demonstrated by the dramatic reduction in the 210 nm negative peak. In terms of the spraying effect, there was a detectable difference in the far-UV CD spectra; however, this difference was observable only at the highest airflow rate through the nozzle (40 L/min).

**Figure 3.**
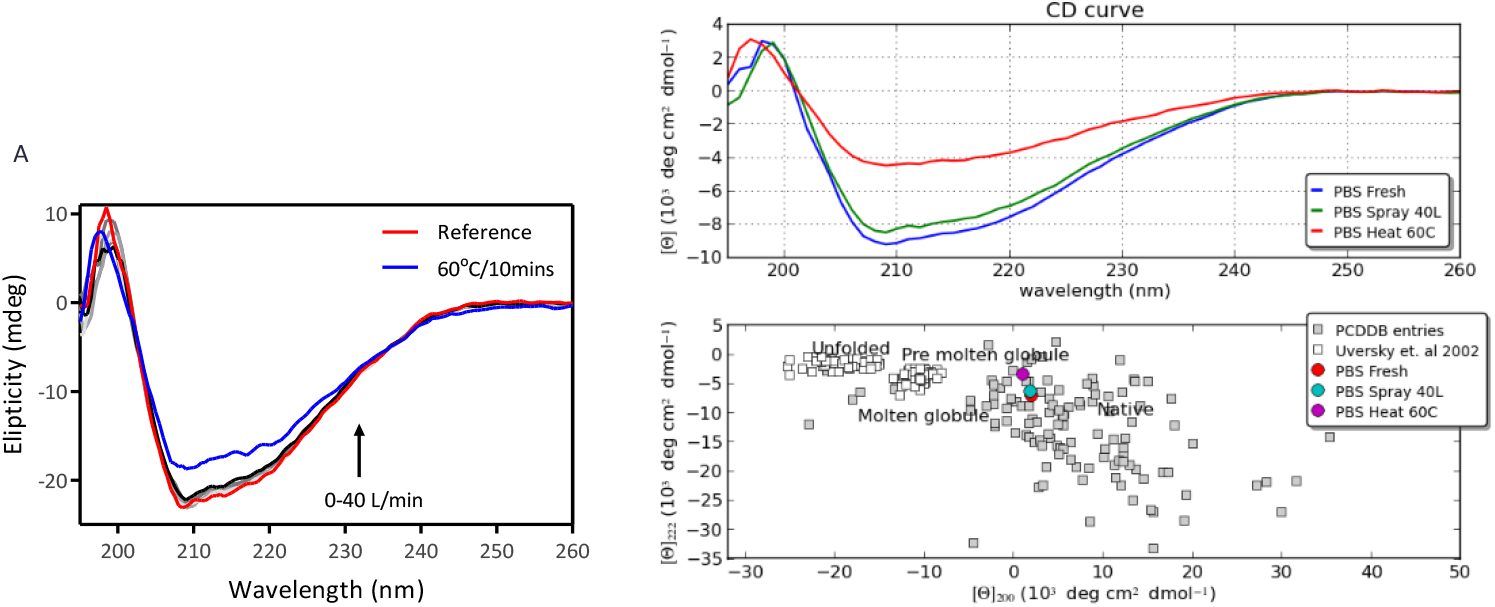
(A) Far-UV CD spectra of lactoferrin in water, atomized at an airflow rate of 0–40 L/min and heated at 60 °C for 10 min. (B) Relative position of lactoferrin samples in CD database with clusters indicating native, molten globule, and unfolded forms.

Spray atomization under the most extreme conditions resulted in only a minimal suppression of the 195 nm peak and a slight decrease in the negative band. Nonetheless, analysis of the data shows that the atomized sample changed from its native population to a pre-molten globule. This indicates that lactoferrin was partially relaxed from its natural folded state by the spray atomization process (Fig. 3B).^2^ In the other samples with milder atomization conditions, the spectra overlap with the original samples. These initial results suggest that the tightly arranged secondary structure of lactoferrin was not disrupted by (a) the mechanical energy required to atomize the protein-containing liquid into droplets or (b) its exposure to the air–water interface. Additionally, the population of the denatured lactoferrin may be too small to elicit a signal detectable by the far-UV CD spectra.

Subsequently, near-UV 250–350 nm CD spectroscopy was utilized to study the tertiary structure of lactoferrin subjected to the effect of spray atomization. In this wavelength region, the primary chromophores are amino acid residues (e.g., Phe, Tyr, Trp) instead of peptide bonds. The near-UV CD spectra of the lactoferrin reference sample showed three distinctive peaks corresponding to the absorption bands of Phe, Tyr, and Trp residues (Fig. 4A). The results from Fig. 4A–B show that as the atomizing airflow rate increased, the 291 nm Trp and 253 nm Phe peaks gradually exhibited a bathochromic red shift of 1–2 nm. The results from Fig. 4A–B suggest that the tertiary structure of lactoferrin was likely disturbed by the spray atomization process, particularly at the spatial arrangement of the Phe, Tyr, and Trp residues. This red shift was observed in both the lactoferrin dissolved in PBS and the lactoferrin dissolved in water, thus indicating a more hydrophobic environment surrounding the Trp and Phe residues.

**Figure 4.**
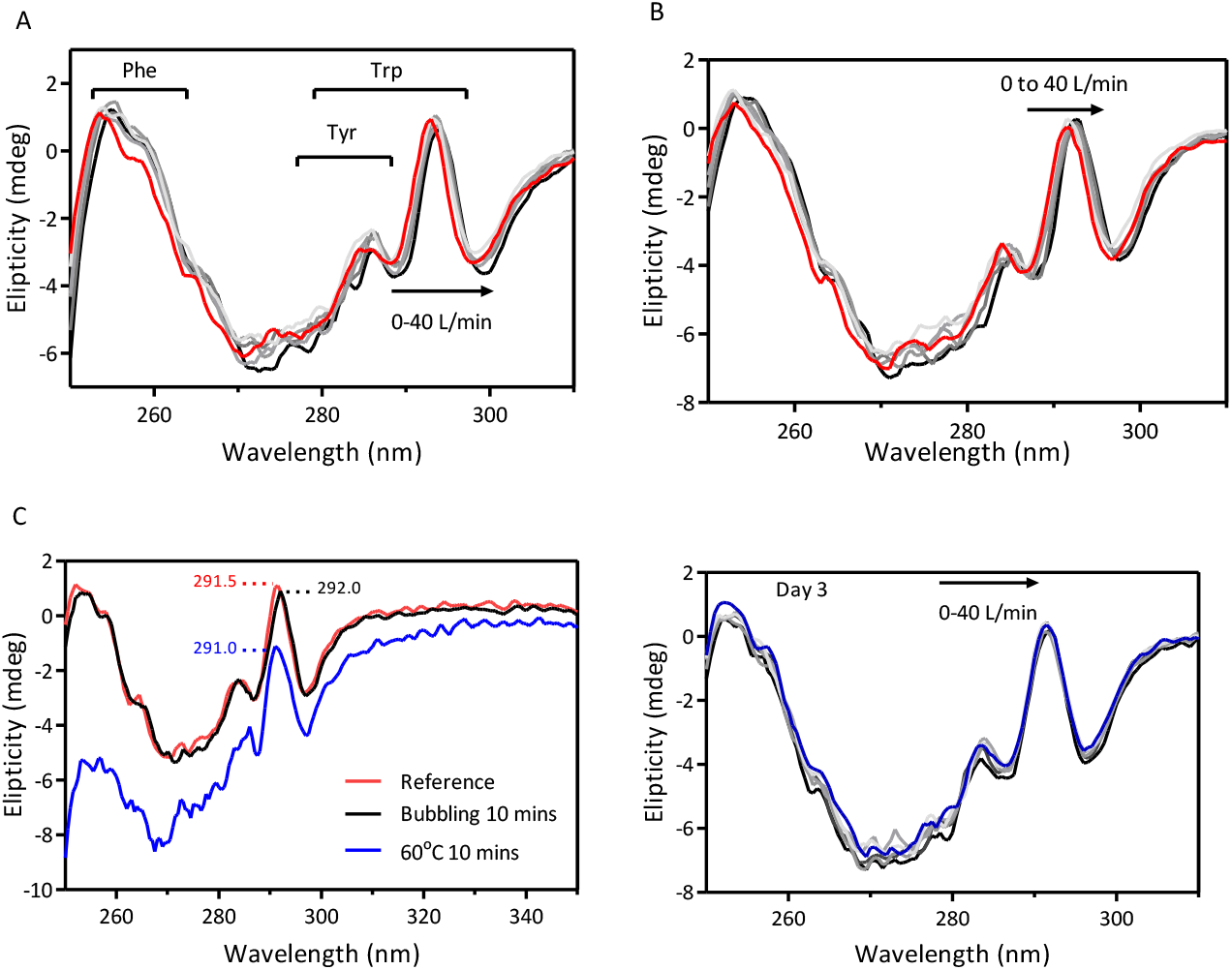
Near-UV CD spectra of lactoferrin in a 0.01 M PBS pH 7.4 solution (A), in water (B), in samples subjected to bubbling with nitrogen gas for 10 min before heating at 60 ° C for 10 min(C); and the near-UV CD spectra of lactoferrin in a 0.01 M PBS pH 7.4 solution at day 3(D).

It is noteworthy that a blue shift is usually observed in chemically induced unfolding studies (Fig. S3C).^48^ The protein is unfolded as the chaotropic agents (e.g., urea, guanidinium chloride) disrupt the hydrogen bonds of the peptide backbone. Subsequently, the aromatic residues (e.g., Phe, Tyr, Trp) that were previously buried inside the hydrophobic pocket and shielded from the aqueous environment are exposed to water.^48^ A blue shift is observed as a result of the electron-deficient environment surrounding these aromatic residues. In contrast, it has been shown that adsorbed lactoferrin molecules at the air–water interface produce a protein film. The top layer of this film is 10–20 Å deep with a high polypeptide volume fraction of 0.5, while the bottom layer is 50–80 Å deep with low polypeptide volume fraction of 0.2.^40^ The polypeptide chain regions in the top dense layer (that is exposed to air) exhibit a strong hydrophobic nature.^40^ As a result, the microenvironment surrounding the aromatic residues Phe and Trp become more hydrophobic, leading to the observed red shift in the 291 nm ellipticity band in the CD spectra.^49^ A similar trend of opposite peak shifting phenomenon was also observed in the case of heating and nitrogen bubbling, thus showcasing different mechanisms of protein denaturation (Fig. 4C).^49^

At low protein concentrations, the disturbance in the spatial arrangement of Trp residues and the associated shift at 291 nm could be reversed through relaxation (Fig. 4D, Fig. 5 and Fig. S3). The results from Fig. S3A–B show that the recorded spectra of all samples at day 3 were practically overlapping as the samples were diluted from operating concentrations of 5–50 mg/mL (employed in SFD^50^ and TFFD^14,26^) to ~1 mg/mL (suitable for CD spectra) and subsequently stored at ambient temperature. In contrast, at higher concentrations (5 mg/mL), lactoferrin molecules irreversibly aggregate into insoluble particles, which is visually detectable is some cases (Fig. 5 and Fig. S3). These results suggest that if the protein concentration is sufficiently high, the exposed aromatic residues of denatured lactoferrin molecules can interact with residues of adjacent molecules to form aggregates. However, the protein can refold into its native state if the protein concentration is low enough to minimize the probability of denatured lactoferrin molecules coming into close proximity.

**Figure 5.**
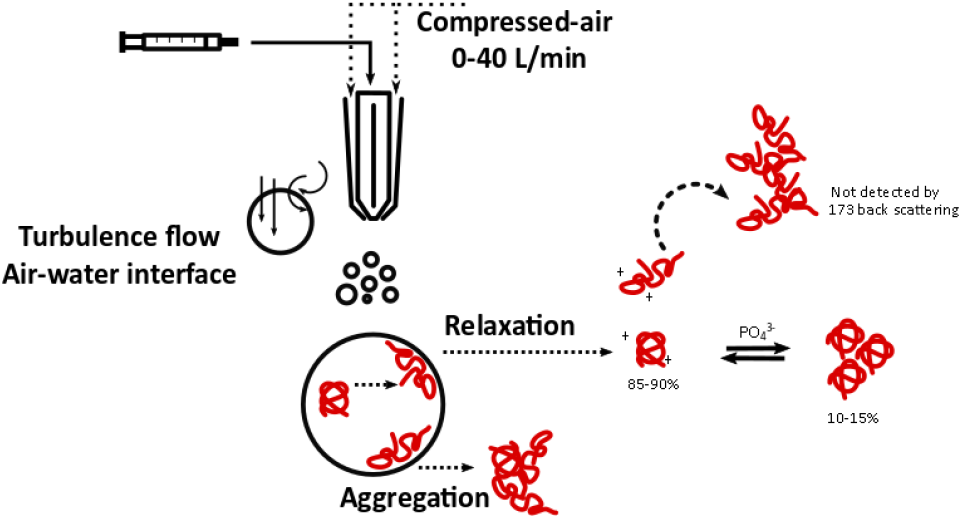
Schematic illustration of the experimental setup for the spray atomization of a lactoferrin solution and the related air–water-induced interfacial denaturation process. The figure illustrates the various protein populations presented in the samples

Dynamic light scattering (DLS) was utilized to characterize the aggregation tendency of lactoferrin samples after spray atomization. DLS volume-based size distribution patterns of freshly prepared lactoferrin reference samples typically consist of two distinctive peaks centered around 10 nm and 50 nm. These peaks can be assigned to the population of protein monomers and oligomers (alternatively known as *high molecular weight* (HMW) species) (Fig. 6A). Any peaks > 1 μm are assigned to aggregates. The area under the curve of the volume distribution of the respective peaks was assigned a relative percentage between the monomers and HMW species. In this regard, lactoferrin in a 0.01 M pH 7.4 PBS solution exhibited about 85–90% monomer content, while the water samples showed 90–99% monomer content (Fig. 6B). These results agree with those obtained by size exclusion chromatography (Fig. 8B). Since lactoferrin is a highly positively charged protein, the negatively charged multivalent phosphate ions can act as physical cross-links to form oligomers via electrostatic attraction.^41^ The protein monomer content was close to 100% in the absence of multivalent ions (Scheme 2 and Fig. 6B).

**Figure 6.**
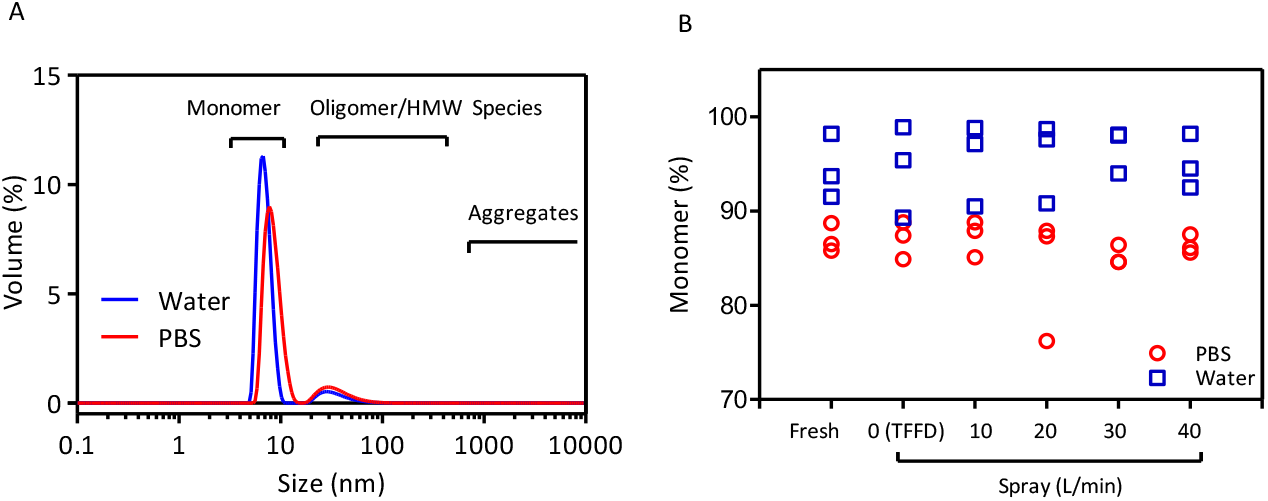
DLS volume distribution of lactoferrin in a 0.01 M PBS pH 7.4 solution and in water (A). The percentage of monomers derived from the area under the curve of peaks center around 5–10 nm (B).

The dual 173° and 13° light scattering angle measurements were used to follow the evolution of small- and large-particle populations, respectively. The derived count rate (or the normalized intensity of scattered light) is a function of the number and size of the particles presented in the field of view.^51^ Although this function is not linear, it can be used to estimate the total number of particles present in the system. Thus, a higher derived count rate value is assigned to a higher number of particles presented in the system, or the particles are significantly larger than the reference.

Immediately after spray atomization, negligible differences between the samples were observed, including the reference, TFFD drop application, and samples atomized at different airflow rates (the black symbols in Fig. 7A). After three days of storage at ambient temperature, the PBS samples at the higher airflow rates showed a slight decrease in count rate from ~25,000 to ~20,000 kcps. This decrease in the small-particle population was concomitant with a dramatic increase in the number of large particles measured using a 13° scattering angle (Fig. 7B). The protein samples that were subjected to spray atomization demonstrated a significant increase in the count rate of large particles: a 20- to 60-fold increase from ~300 kcps to ~13,000 kcps and ~65,000 kcps, respectively (Fig. 7B). These changes were also accompanied by a 10-fold increase in the z-average value (Fig. 7C). It is also noteworthy that even in the absence of atomizing air (TFFD, 0 L/min), the mere act of dispensing the protein solution through the two-fluid nozzle also affected the aggregation tendency of lactoferrin. Fig. 7B shows that at day 3, the count rate of reference and the count rate of the TFFD sample were 323 kcps and 833 kcps, respectively. Nonetheless, dispensing the protein solution (thus exposing the protein to shear stress) had a minimal effect on the drop application of TFFD compared to the detrimental effect of high-shear spray atomization.

**Figure 7.**
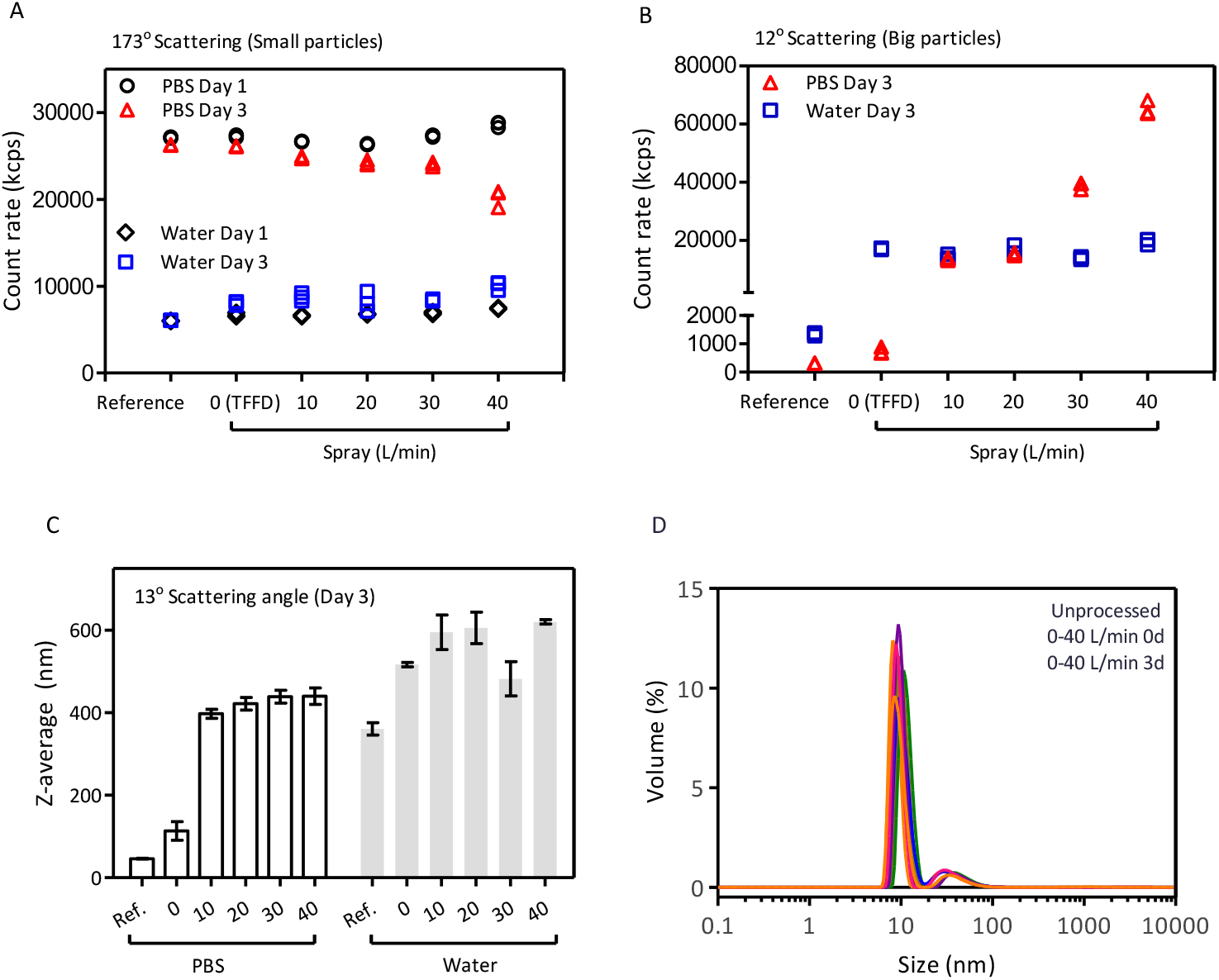
Derived count rate of lactoferrin in water and in PBS at day 1 and day 3 after spray atomization, measured using a 173° scattering angle (A) and a 13° scattering angle (B) to monitor small- and large-particle populations, respectively. The z-average of lactoferrin in water and PBS samples at day 3 (C). Volume-based particle size distribution of various reference and atomized lactoferrin samples (D).

Surprisingly, there was no detectable difference in the 13° scattering angle measurements among the protein-in-water samples, even though protein aggregation in these samples was visually evident (Fig. 7A–B and Fig. S4). These results suggest that the visually observed protein aggregates were larger than the DLS limit of detection, thus they were not added to the derived count rate. Another interesting point is that the relative percentage of monomer and oligomer peaks remained relatively constant, even in samples with visually confirmed aggregation (Fig. 7D). The percentage of the area under the curve of monomer and oligomer peaks remained at 85–90% and 10–15%, respectively. This observation suggests that lactoferrin monomers and oligomers are in an equilibrium dependent on the pH level and the concentration of phosphate ions (Fig. 5). As the protein monomers were denatured by spray atomization, the population of denatured monomers aggregated into large insoluble particles. Subsequently, the loosely connected oligomers were disassembled to maintain the equilibrium. Consequently, only negligible changes were observed in the relative ratios of the area under the curve of monomer and oligomer peaks in the DLS volume-based distribution. On the other hand, the absolute value of the number of monomers was expected to decrease, with a corresponding increase in oligomers. Later in this study, size exclusion chromatography is used to corroborate this assumption (Fig. 8A–B).

**Figure 8.**
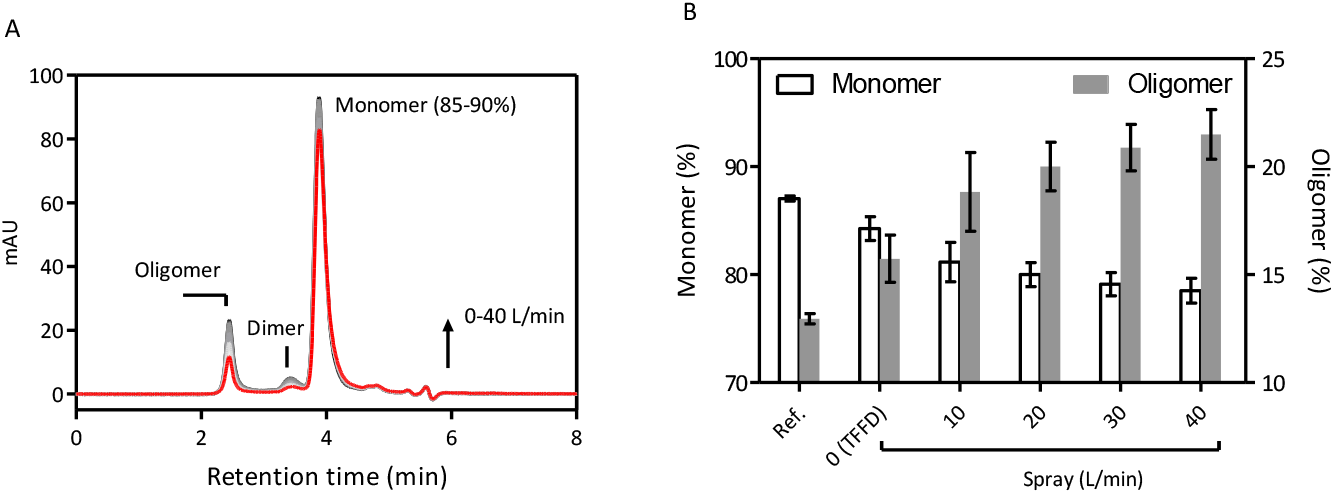
Unprocessed and processed LF solutions: SEC chromatogram (A) and percentage of monomer and oligomer content (B).

In summary, DLS is a useful tool for screening protein solution samples based on their protein aggregation tendencies. DLS provides fast, non-invasive, qualitative, and relatively accurate characterizations of the protein population as a whole.^52^ The protein samples are complex systems with many components such as monomers, oligomers, soluble and insoluble aggregates, and even the dust particles inadvertently introduced during the preparation and handling phase. Each of these components interact in the interplay and equilibrium. Hence, a combination of synergistic parameters was used to provide both quantitative and qualitative analyses of the protein samples. Specifically, the lactoferrin aqueous samples in this study were characterized using different viewing perspectives, including the z-average, the volume distribution, forward 13° and backward 173° scattering, and—most importantly, the derived count rate.

In addition, SEC was employed to determine the amount of monomers and oligomers presented in the sample. The upper detection limit of the column used in this study is 1,250 kDa, corresponding to the aggregation of 15 lactoferrin molecules. Indeed, the SEC chromatogram agrees with the data from DLS characterization, in which the monomer content in the reference samples comprised 85–90% of the protein population (Fig. 8A–B). The remaining 10–15% was attributed to protein oligomers.^3,53^ Upon spray-atomizing at a 0 L/min (TFFD relevant condition) and a 10–40 L/min airflow rate, the monomer content gradually decreased, accompanied by an increase in oligomer content (Fig. 8B). It is notable that sodium dodecyl sulphate–polyacrylamide gel electrophoresis (SDS-PAGE), did not differentiate freshly prepared, thermally, and physically denatured protein samples with visual denaturation cues (Data not shown). This result was possibly due to the low upper molecular weight limit of 250 kDa of the reference protein ladder, which translates into a 3-lactoferrin-molecule aggregate.

The aggregation tendency of the lactoferrin solution subjected to spray atomization was further characterized using MFI, with an emphasis on the morphology of the aggregates (Fig. 9A–B). Fig. 9B shows several heat maps of the individual particles detected by the MFI. The morphology of individual particles can be estimated depending on their relative position. Specifically, the data points in the upper-right quadrant (1,1) represent circular particles, while data points in the lower-left quadrant (0,0) represent elongated, irregularly shaped particles. The protein particles were also differentiated based on their ECD, with smaller ECD particles represented by warmer colors and larger ECD particles represented by cooler colors. Additionally, protein aggregates are rarely perfectly circular in shape, while microbubbles and silicon oil droplets are usually circular; therefore, only data points with circularity < 0.9 and aspect ratio < 0.9 are presented.

**Figure 9.**
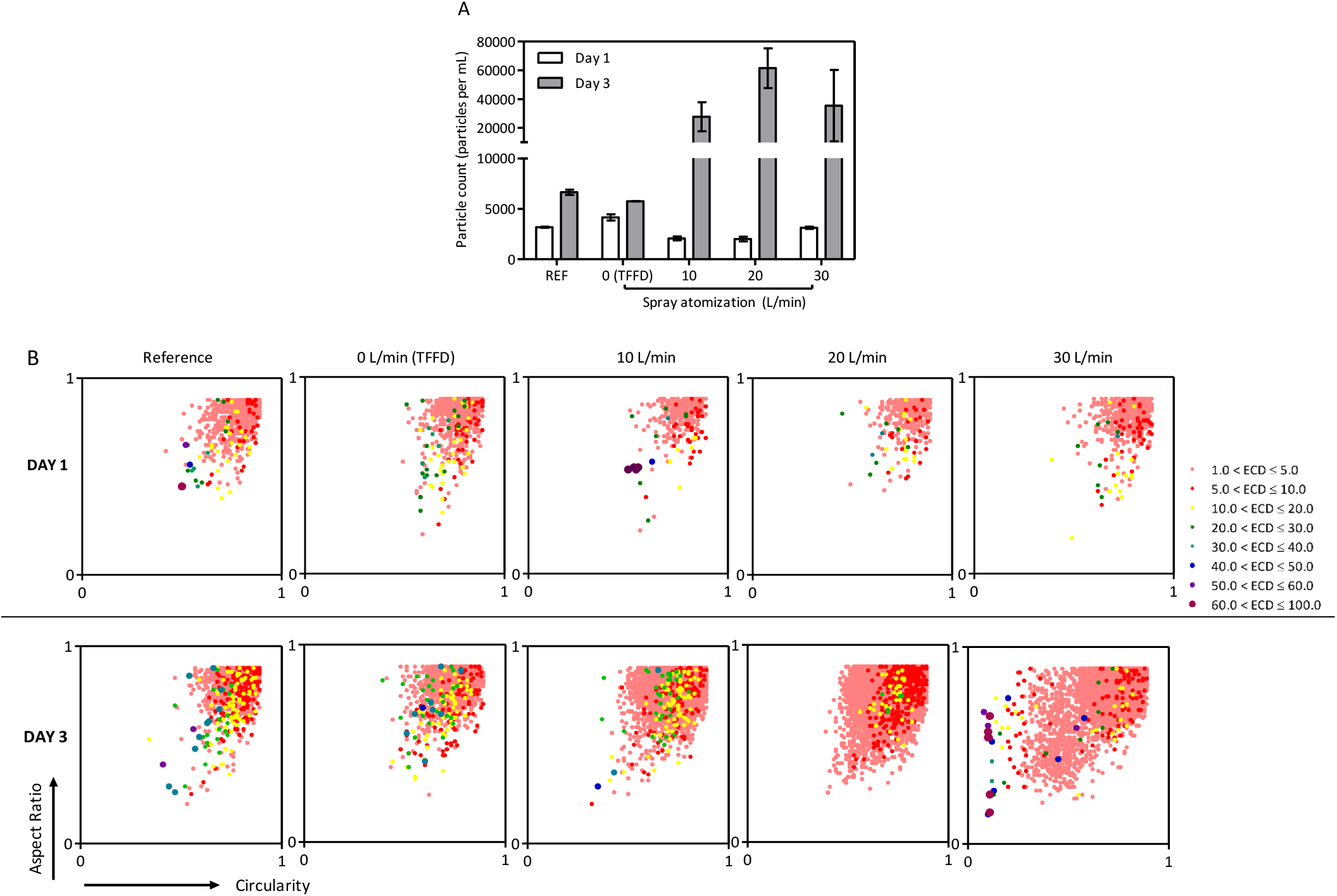
Total particle counts (1–300 μm) in spray-atomized lactoferrin samples (A). Scatter plot of individual aggregates detected by MFI in 1.0 mg/mL lactoferrin solution in 0.01M pH 7.4 PBS (B).

Initially after spray atomization, there are only statistically insignificant differences in particle counts between the reference and processed samples (Fig. 9A–B). The particle counts even decrease, from around 3,000 in the reference sample to ~2,000 in the spray-atomized samples. However, at day 3, a significant increase in particles can be observed in all spray-atomized samples compared to the reference and TFFD drop application samples. This result suggests that shear stress might break up some intrinsic lactoferrin aggregates into smaller particles below the detection limit of MFI (1 μm), thus reducing the particle counts. However, with the progression of time, the denatured lactoferrin molecules can aggregate into larger particles and thus significantly increase the particle counts shown in both the bar graph and heat map (Fig. 9A–B).

The heat maps in Fig. 9B show that the aggregates in the spray-atomized sample (10–30 L/min) exhibited changes both in size and morphology. The distribution of particles in the reference and TFFD samples was confined to the upper-right quadrant, while they gradually expand to the lower-left quadrant. This indicates the formation of a population of more elongated aggregates. Additionally, the sample atomized at 30 L/min also exhibited the widest particle size distribution and morphology, suggesting a relatively heterogeneous nature. Data from Fig. 9A–B suggest that spray atomizing does not simply increase the number of protein aggregates; it also alters the morphology of the aggregates, forming larger and irregularly shaped particles.

The denatured, and subsequently aggregated, lactoferrin solutions were further investigated to determine their surface adsorption tendencies via a drop-shape-analysis tensiometer. Immediately after atomizing the liquid through two-fluid nozzles into small droplets, subtle differences in the adsorption kinetic curve were detected (Fig. 10A). Specifically, the equilibrium surface tension at 600 s demonstrated a decreasing trend as the airflow rate in the atomizing nozzle was increased (Fig. 10C). With the exception of the TFFD-relevant 0 L/min processing condition for drop application, the other three samples with SFD-relevant atomizing airflow rates of 10–40 L/min exhibited a lower equilibrium interfacial tension than the reference sample.

**Figure 10.**
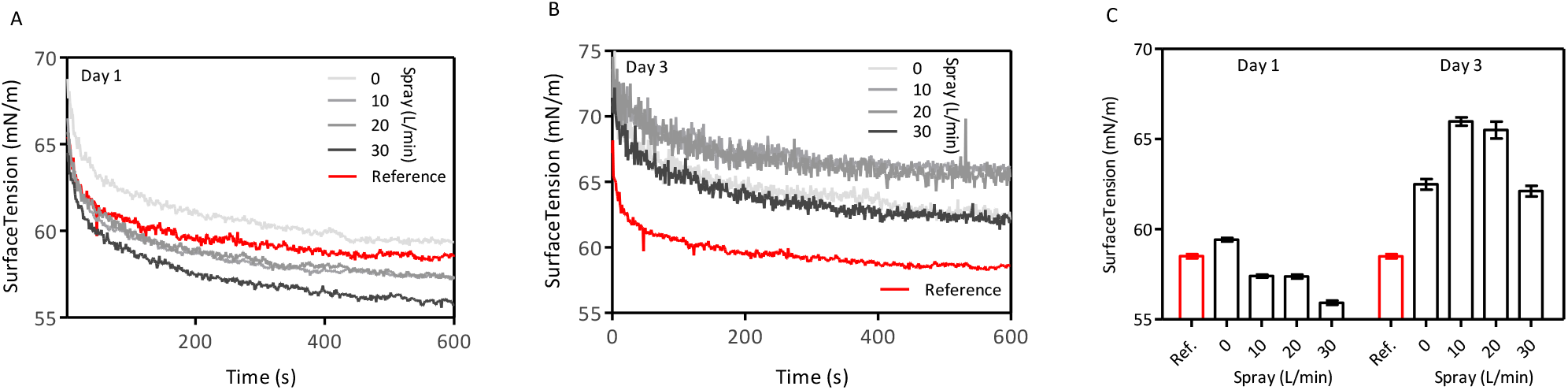
Surface tension of the air–water interfaces of lactoferrin reference and spray-atomized solutions at day 1(A), day 3 (B), and the average surface tension over the last 50 s of measurement (C).

Interestingly, DLS confirmed that this reduction in surface tension was reversed as the aggregation proceeded (Fig. 10B–C). All SFD-relevant spray-atomized protein solution samples exhibited higher equilibrium interfacial tension than the reference. These results, along with previous findings, suggest that spray-induced conformational changes in lactoferrin molecules might expose the protein’s hydrophobic core. As a result, these partially unfolded proteins have a stronger tendency to adsorb onto the air–water interface, thus decreasing the overall surface tension of the pendant droplet (Fig. 10A).^13^ However, the presence of large micron-sized aggregates will adversely affect the surface activity, resulting in higher surface tension. It is suggested that protein aggregation can produce a weaker interfacial network than unprocessed protein samples because fewer proteins are available to interact at the interface.

Lactoferrin solutions were also spray-atomized from various heights to examine the effect of air–water interface exposure time. The underlying assumption in this set of experiments is that, given the same surface-to-volume ratio, the duration of a protein’s exposure to the hydrophobic environment correlates with the ratio of denaturation. However, this effect could not be detected within the time range relevant to the SFD setup in this study, which is typically < 1 s (Fig. 12A–C). It was suggested that the denaturation or unfolding of proteins via exposure to the air–water interface, although reversible, usually happens within 10 ns.^4^ Initial adsorption is usually controlled by diffusion, but further adsorption requires the protein to overcome an energy barrier. Hence, protein migration time is much slower than the denaturation step, which happens nearly instantaneously after the atomization of the droplet and the associated formation of new surface areas (Fig. 11).

**Figure 11.**
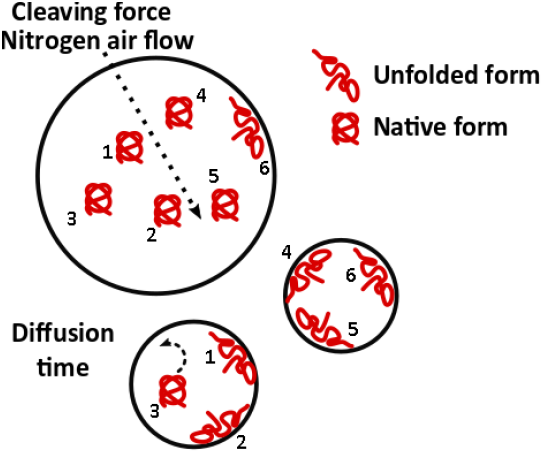
Proposed schematic illustration of protein diffusion and denaturation time during process diminution and flying time, where diffusion time is suggested to be significantly longer than the flying time.

**Figure 12.**
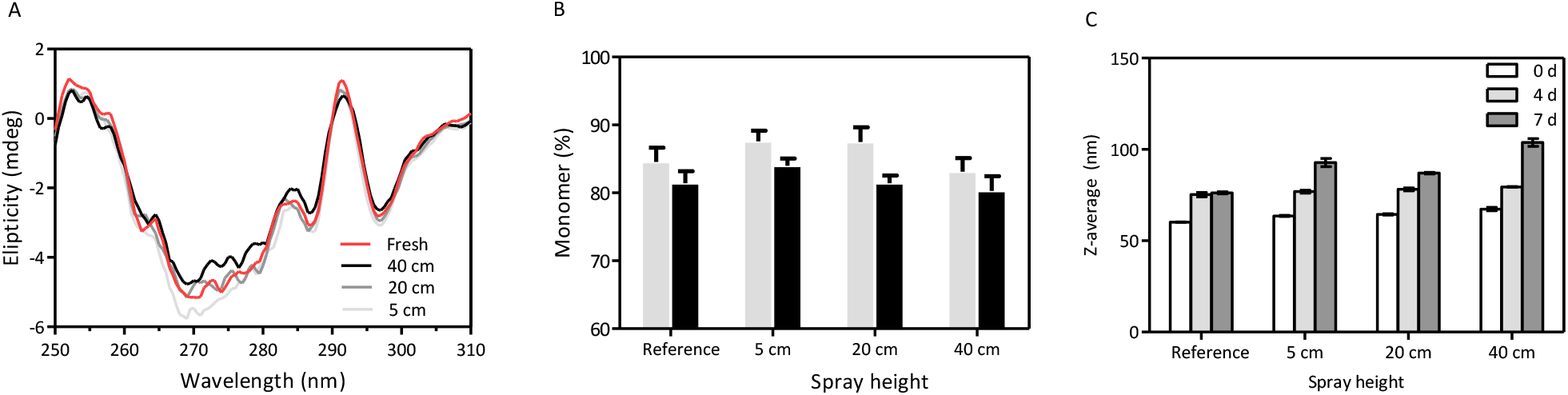
Near-UV CD spectra (A), SEC monomer content (B), and z-average values (C) of lactoferrin samples sprayed at different heights ranging from 5–40 cm from the top of the nozzle to the bottom of the collecting vessel.

The result from Fig. 12A shows that the near-UV CD spectra of samples with varied spray heights practically overlapped, and no significant shift in peak position at 290 nm was observed. There was only a slight perturbance in the 270– 285 nm region. Additionally, the percentage of monomer content from SEC (Fig. 12B) and aggregation tendency obtained via DLS (Fig. 12C) all suggest there is no significant difference.

Taken altogether, these results suggest that the diffusion of proteins to the interface within the time scale of the spraying process contributes only a small portion of the overall detrimental effect on protein stability. More conformational changes occur relatively quickly and overshadow the effect of diffusion-based denaturation. To gain a rough approximation of the diffusion time of lactoferrin from bulk liquid to the air–liquid interface, laser diffraction was used to examine the droplet diameters that correspond to the various spray atomizing conditions. The results (presented in Fig. 13) demonstrate a sharp reduction in the mean droplet diameter, from ~200 μm at an airflow rate of 10 L/min to 50 μm at a 20 L/min airflow rate. At airflow rates above 20 L/min, the droplet diameter showed diminishing returns and stabilized at about 20–30 μm.

**Figure 13.**
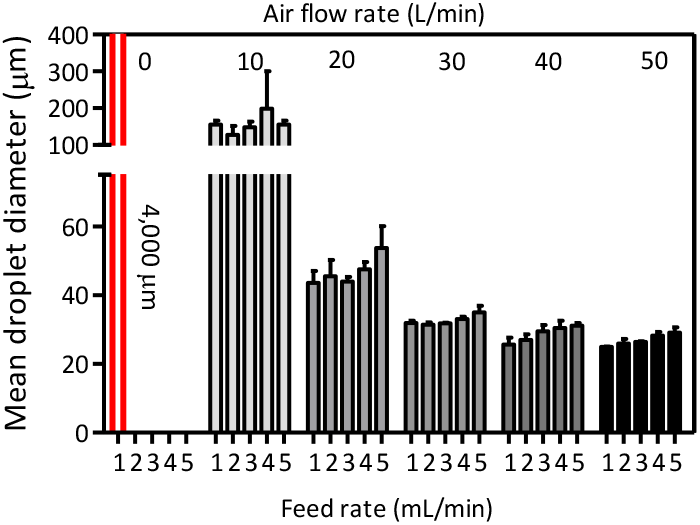
Mean droplet diameter of spray-atomized liquid samples at varying airflow rates and feed rates.

The drop diameter is about 4.0 mm or 4,000 μm for liquid passing through the nozzle without airflow, equivalent to TFFD. The time scale (*t*) required for the protein to traverse distance R in three-dimensional space with a diffusion coefficient (D) was estimated using the simplified Fick’s second law for Brownian particle and mean squared displacement: *t* = R^2^/6D. ^54^ Specifically, the diffusion coefficient of lactoferrin in water obtained by DLS was 11–12 μm^2^/s. The protein is assumed to have reached the surface if the traversed distance equals the radius of the spray-atomized droplet (R). Considering this, the results from Table 1 show that for the droplets with a 2,000 μm radius in the TFFD processing technique, it is nearly impossible for the protein to effectively diffuse from the bulk liquid within the large drop to the interface in order to cause any significant detectable unfolding. Regarding the SFD atomizing process, this estimation demonstrates that, within the scope of the spray parameters set in this study, the protein diffusion time falls in the range of 1–100 s. Meanwhile, the sprayed droplet travel time (before its impact with the vessel wall) falls in the microsecond range. Therefore, it is reasonable that a change in spraying height did not result in detectable changes in the conformation and aggregation tendency of lactoferrin (Fig. 12A–C). Altogether, the results suggest that most of the lactoferrin denaturation occurs almost immediately upon formation of a new interfacial area by spray atomization to form micron-sized droplets with protein molecules located sufficiently close to the newly formed liquid–air interface (Fig. 11, proteins 1, 2, 4, and 5). Prolonging the exposure of droplets in the air by a fraction of a second had no significant impact because protein diffusion is relatively slow (Fig. 11, protein 3).

**Table 1.** Estimated time required for lactoferrin to diffuse in spray-atomized droplets.

| Droplet radius ($\mu\text{m}$) | | Diffuse time bulk-interface |
| --- | --- | --- |
| 2,000 $\mu\text{m}$ (TFFD) | | 16.8 h |
| 100 $\mu\text{m}$ | (SFD) | 150 s |
| 25 $\mu\text{m}$ | | 9 s |
| 15 $\mu\text{m}$ | | 3 s |
| 10 $\mu\text{m}$ | | 1 s |

## DISCUSSION

### Implications of stresses incurred during droplet formation on lactoferrin conformation

This study compares the droplet formation process of SFD and TFFD, two powder-processing techniques used for producing inhalable dry powders. It has been reported that shear stress arising from the pumping, filtering, and spraying process can affect protein stability.^55^ Subjecting the dissolved proteins to a steep velocity gradient in the two-fluid nozzle can cause the polypeptide chain to unfold and subsequently aggregate.^8,28,30^ However, several studies have reported that a controlled linear constant shear velocity gradient (e.g., the gradient encountered during pumping and filtering) is insufficient to cause denaturation of globular proteins.^8,28,30^ The authors conclude that even an extraordinarily high shear rate of 10^7^ s^−1^ is insufficient to significantly denature cytochrome c, a small globular protein.^30^ Similarly, in the absence of air, rhGH and rhDNase remained relatively intact even under extreme shear.^5^ In contrast, both proteins exhibited a significant increase in aggregation tendency in the presence of air at air–water interfaces. These findings suggest that exposure to air– water interfacial areas is the crucial factor in protein denaturation.^22^ In light of this, powder processing technologies that require spray atomization (e.g., spray drying, SFD,^50^ and spray freezing into liquid^56,57^) inherently reduce the stability of macromolecules due to the high surface area–to-volume ratio of the resultant droplets.

Several studies have observed that the two-fluid nozzle used in spray drying eliminated as much as 23% of the activity of lactate dehydrogenase; and atomization, dehydration, and circulation contributed equally to this problem.^28^ Engstrom compared LDH activities and also concluded that TFFD can maintain higher LDH activity than SFD.^15–17,58^ The present study complements this work by providing a mechanistic understanding of protein denaturation.

Concerning the susceptibility of proteins to denaturation induced by air–water interfaces, the rate of denaturation depends on the surface-active properties of the protein. Specifically, the tendency to adsorb onto the interface (e.g., greater foaming ability) increases the level of denaturation.^31^ In this aspect, the equilibrium surface tension of lactoferrin (or the surface activity of lactoferrin) is similar to iGg,^43^ rhGH,^59^ mAbs,^60^ and lysozyme^61^ (with an equilibrium surface tension of 20 μM ~ 55–60 mN/m). These are proteins known to degrade with exposure to the air–water interface through the aeration of the solution.

Indeed, this study demonstrates that lactoferrin underwent structural changes upon exposure to a large air–water interface in the micron-sized atomized droplets required by the SFD process. The diameter of the spray-atomized droplets in this study (20–150 μm) is relevant, and in certain cases the droplets are even larger than those observed in previously reported studies.^62,63^ Thus, droplet diameter is a good indication of whether mitigating approaches should be applied to prevent protein denaturation.

Finally, this study confirms that technologies that require spray-atomizing to prepare inhalable powders (e.g., SFD and spray drying) have an inherent detrimental effect on surface-active therapeutic proteins, and possibly other biological macromolecules. Considering this, TFFD offers much milder overall processing conditions because of its low-shear stress and its use of drops that have a very low surface area–to-volume ratio. This makes TFFD more suitable for processing therapeutic macromolecules while minimizing their denaturation.

## CONCLUSION

This study shows that lactoferrin was denatured by spray atomization processes that form micron-sized liquid droplets with high surface area–to-volume ratios, and it exhibited fewer protein monomers as well as an altered tertiary structure. The primary mechanism of denaturation was exposure of the protein to the expansive air–water interface that results from the formation of micron-sized droplets. Subsequently, these degraded protein molecules can aggregate into insoluble particles detectable by 173° forward-angle DLS. In the context of the stresses encountered during the droplet formation stage of the TFFD process, the utilization of millimeter-sized (i.e., larger) drops with low shear used in the TFFD process greatly improved protein stability.

## ASSOCIATED CONTENT

### Supporting Information

This material is available free of charge via the Internet at http://pubs.acs.org.

## AUTHOR INFORMATION

### Author Contributions

The manuscript was written through contributions of all authors. / All authors have given approval to the final version of the manuscript.

## ABBREVIATIONS

TFFD: Thin-film-freeze-drying
SFD: Spray-freeze-drying
SEC: Size Exclusion Chromatography
CD: Circular dichroism

