## Supplemental Supporting Information for "Degradation of lactoferrin caused by droplet atomization process via two-fluid nozzle: The detrimental effect of air–water interfaces"

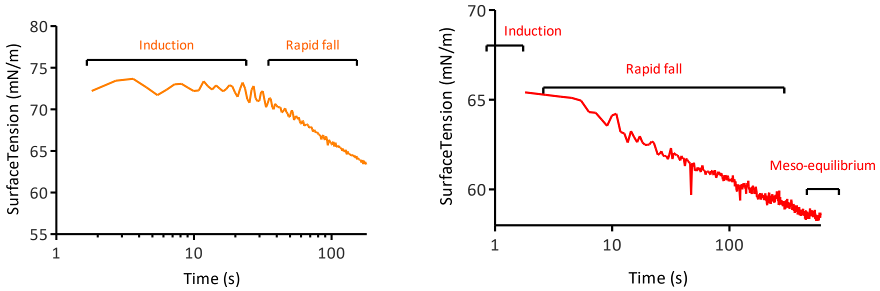


**Figure S1**. Representative curve of dynamic surface tension experiment with surface-active molecules (lactoferrin).

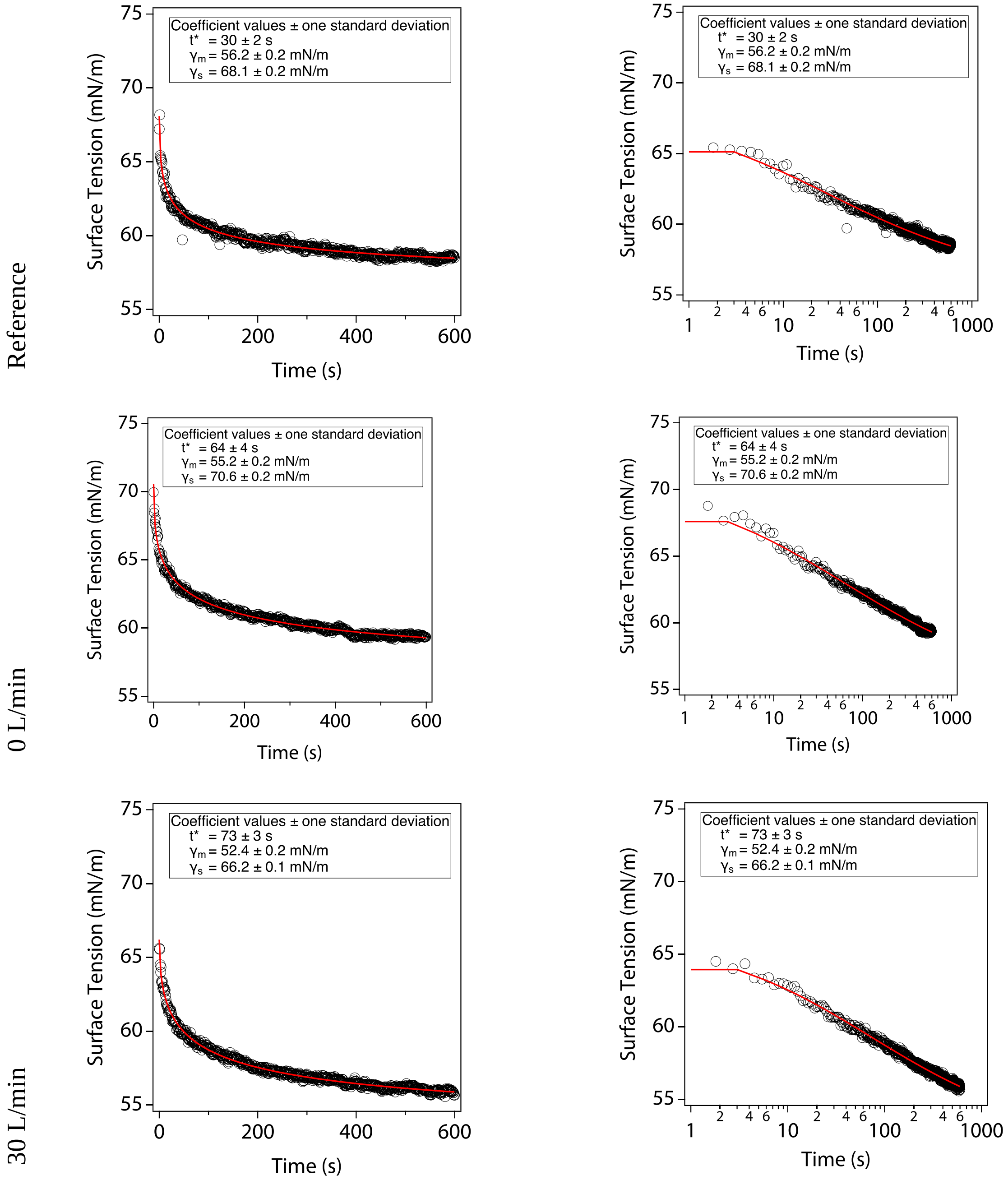


**Figure S2**. Obtaining $t^*$ time by fitting the dynamic surface tension curve of lactoferrin (in 0.01 M PBS pH 7.4) to the Hua–Rosen model.

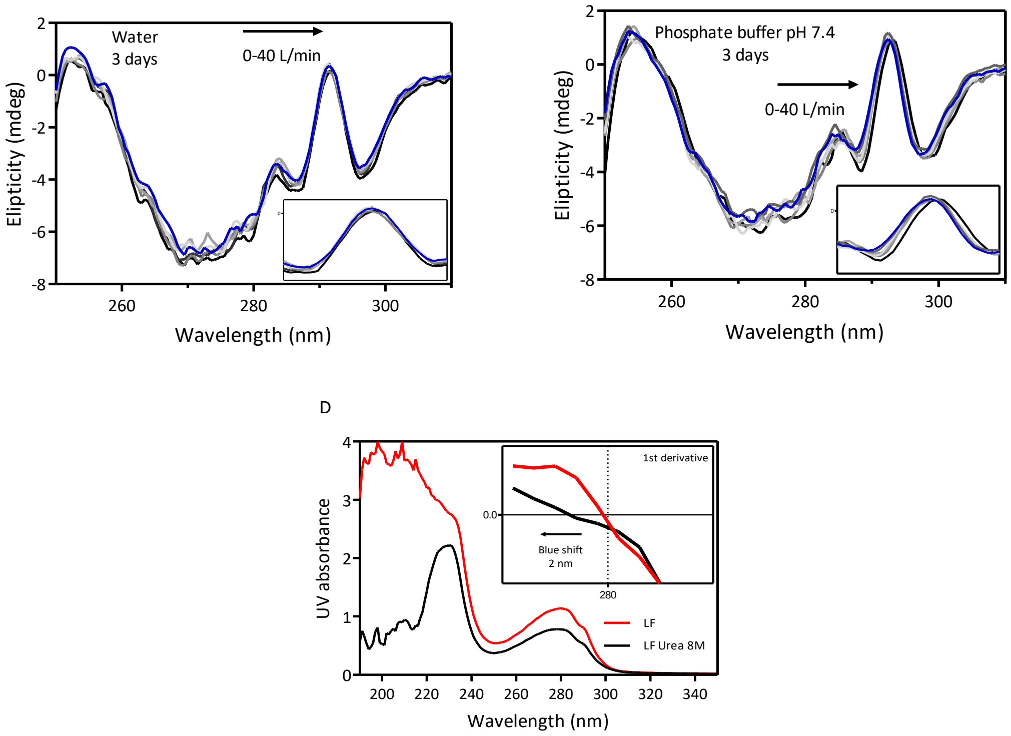


**Figure S3**. Near-UV CD spectra of lactoferrin solutions in water (A) and in 0.01 M PBS pH 7.4 (B) after 3-day storage at ambient temperature.

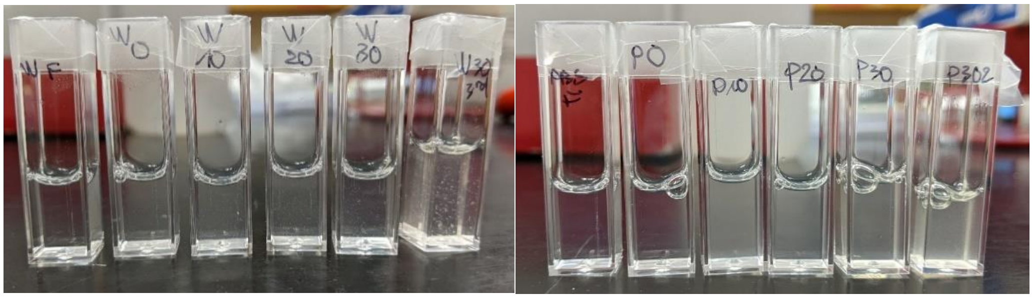

**Figure S4**. Photos of 5 mg/mL lactoferrin solutions in water and in 0.01 M PBS pH 7.4, unprocessed and sprayed with a 0–40 L/min airflow rate.

Graphic entry for the Table of Contents (TOC)

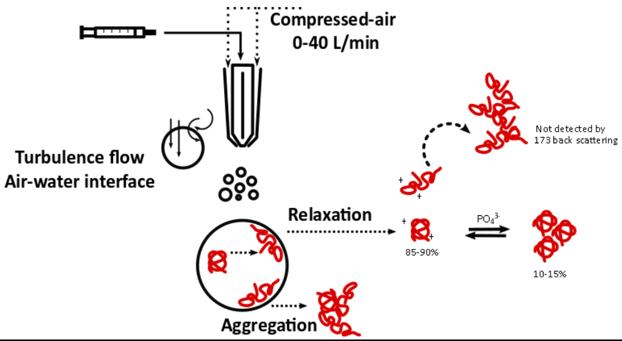
